# PBRM-1/PBAF-regulated genes in a multipotent progenitor

**DOI:** 10.1101/2023.10.16.562537

**Authors:** Laura D. Mathies, Andrew C. Kim, Evan M. Soukup, Alan’da E. Thomas, Jill C. Bettinger

## Abstract

The *Caenorhabditis elegans* somatic gonadal precursors (SGPs) are multipotent progenitors that generate all somatic cells of the adult reproductive system. The two SGPs originate in the mesodermal layer and are born through a division that produces one SGP and one head mesodermal cell (hmc). One hmc terminally differentiates and the other dies by programmed cell death. The PBAF chromatin remodeling complex promotes the multipotent SGP fate. Complete loss of PBAF causes lethality, so we used a combination of Cre/lox recombination and GFP nanobody-directed protein degradation to eliminate PBRM-1, the signature subunit of the PBAF complex, from 83 mesodermal cells, including SGPs, body muscles, and the hmc. We used RNA sequencing to identify genes acting downstream of PBAF in these cells and identified 1955 transcripts that were significantly differentially expressed between *pbrm-1(-)* and *pbrm-1(+)* in the mesoderm of L1 larvae. We found that genes involved in muscle cell function were overrepresented; most of these genes had lower expression in the absence of PBRM-1, suggesting that PBAF promotes muscle differentiation. Among the differentially expressed genes were 125 genes that are normally expressed at higher levels in SGP *vs*. hmc and positively regulated by *pbrm-1* and 53 that are normally expressed at higher levels in hmc *vs*. SGP and are negatively regulated by *pbrm-1;* these are candidate regulators of the SGP/hmc fate decision. We validated one candidate gene using a fluorescent reporter; the *hsp-12.3* reporter was derepressed in SGPs in *pbrm-*1 mutants, suggesting that *hsp-12.3* expression is normally repressed by *pbrm-1* in SGPs.

## Introduction

SWItching defective/Sucrose Non-Fermenting (SWI/SNF) chromatin remodeling complexes are multi-protein assemblies that regulate gene expression by altering chromatin structure (Clapier and Cairns 2009). Two major classes of SWI/SNF complexes are BAF (BRM/BRG-associated factors) and PBAF (Polybromo-associated BAF) (reviewed in Wu *et al*. 2009). In mammals, molecularly distinct BAF complexes are found in pluripotent embryonic stem cells (esBAF), multipotent neural progenitors (npBAF), and differentiated neurons (nBAF) (Lessard *et al*. 2007; Ho *et al*. 2009; Lessard and Crabtree 2010). SWI/SNF subunits facilitate the reprogramming of differentiated cells into pluripotent stem cells (Singhal *et al*. 2010), underscoring the importance of SWI/SNF complexes in the regulation of cellular potential.

The *C. elegans* somatic gonadal precursors (SGPs) are multipotent progenitors that generate all 143 cells of the somatic gonad. The two SGPs, Z1 and Z4, are born from cell divisions that produce one SGP and one head mesodermal cell (hmc) (Sulston *et al*. 1983). After their births, the SGPs migrate posteriorly and coalesce with the two primordial germ cells (PGCs) to form the gonad primordium. Each SGP generates one of the two U-shaped arms of the adult gonad (Kimble and Hirsh 1979). The hmcs migrate anteriorly where one cell dies by programmed cell death and the other differentiates as the single hmc. We previously defined the transcriptomes of sorted SGPs and hmcs and found that they had numerous transcriptional differences: ∼3000 genes had higher expression in SGP than hmc (SGP-biased), and a similar number had higher expression in hmc than SGP (hmc-biased) (Mathies *et al*. 2019). Thus, two sister cells that were produced by a single cell division have very different gene expression profiles consistent with their different cellular potentials (terminally differentiated *vs.* multipotent).

Little is known about the regulation of the SGP/hmc fate decision. Mutations in four genes, the conserved mesoderm regulator *hnd-1/*dHand and three genes encoding subunits of the PBAF chromatin remodeling complex, play a role in regulating this cell fate decision (Large and Mathies 2014). Animals carrying mutations in any of the genes have two incompletely penetrant phenotypes: 1) SGPs can be absent from the gonad primordium resulting in adults with missing gonad arms and 2) SGPs can have gene expression patterns characteristic of both SGP and hmc. Both phenotypes can be explained by the partial transformation of SGPs into hmcs and suggest that these genes are important for distinguishing multipotent SGPs from their differentiated hmc sisters. The involvement of three PBAF genes, including the signature subunit *pbrm-1/*Polybromo and the core ATPase *swsn-4,* strongly suggests that *C. elegans* PBAF regulates cellular potential, as has been shown for mammalian BAF complexes.

Here, we used a combination of Cre/lox recombination and GFP nanobody-directed protein degradation to deplete *pbrm-1* mRNA and protein from mesodermal tissues, including SGPs. Mesodermal inactivation of *pbrm-1* resulted in a strong loss-of-function phenotype in the somatic gonad, without the high degree of lethality associated with null alleles of the gene. Using this conditional inactivation strategy, we identified 1955 genes that were differentially expressed in the mesoderm between *pbrm-1(-)* and *pbrm-1(+)*. Genes implicated in muscle function were overrepresented among the differentially expressed genes (DEGs), suggesting a role for PBRM-1 in muscle cell differentiation. To find genes that may be important for the SGP/hmc fate decision, we utilized our existing gene expression dataset from sorted SGPs and hmcs (Mathies *et al*. 2019) to identify 178 candidate mediators of the *pbrm-1* effect on the SGP/hmc fate decision. We used a fluorescent reporter to validate one candidate gene, *hsp-12.3,* which had hmc-biased expression in wild-type animals and became derepressed in SGPs of *pbrm-1* mutants, suggesting that *pbrm-1’s* repression of *hsp-12.3* is an aspect of the SGP fate.

## Materials and Methods

### Strains

*C. elegans* strains were cultured as described previously (Brenner 1974; Wood 1988). All strains were grown at 20°C unless otherwise specified and were derived from the Bristol strain N2. Strains were obtained from the *Caenorhabditis* Genetics Center or were generated as described below. A complete list of strains is included in Supplemental Material (File S1, Table S1). The following strains were generated for this study:

RA650 pbrm-1(rd29[pbrm-1::GFP::3xFlag])
RA661 *pbrm-1(rd31[pbrm-1::GFP(flox)])*
RA663 *pbrm-1(rd31[pbrm-1::GFP(flox)]); rdIs67[hnd-1p::CRE + unc-119(+)]*
RA678 *pbrm-1(rd31[pbrm-1::GFP(flox)]); rdIs74[hnd-1p::GFP-nanobody::ZIF-1::Cre + unc-119(+)]*
RA685 *pbrm-1(rd31); rdIs79[hnd-1p::ZIF-1 + unc-119(+)]*

### Generation of loxP-flanked *pbrm-1::GFP*

GFP was inserted just before the *pbrm-1* stop codon using CRISPR/Cas9 genome editing. A repair plasmid was generated by cloning sequences flanking the intended insertion site (homology arms) into pDD282 (Addgene #66823) digested with AvrII and SpeI. The left homology arm was amplified by PCR using primers RA1386 and RA1387 and the right homology arm was synthesized as a gblock gene fragment (IDT, Skokie, Illinois). The guide sequence was cloned into PU6::unc-119_sgRNA (Addgene #46169) (Friedland *et al*. 2013) using the Q5 site directed mutagenesis kit (NEB, Ipswich, MA) and primers RA1318 and RA1063. Insertions were created following a published protocol (Dickinson *et al*. 2015). Briefly, repair (10 ng/μl), P*eft-3::*Cas9 (50 ng/μl), and guide (50 ng/μl) plasmids were injected into N2 worms with fluorescent co-injection markers. Insertions were selected using a combination of hygromycin resistance and the *sqt-1(d)* roller phenotype; both markers are contained within the self-excising selection cassette of pDD282. The selection cassette was removed by heat shock to create RA650, which retains one loxP site in the final intron between GFP and 3xFlag (Fig. 1A). A second loxP site was inserted 721 bp upstream of the start of *pbrm-1b* using CRISPR/Cas9. The guide sequence was cloned into PU6::unc-119_sgRNA (Addgene #46169) (Friedland *et al*. 2013) using primers RA1063 and RA1413. The repair template contained 36 nucleotide homology arms flanking the loxP sequence and was synthesized as an Ultramer DNA oligo (IDT, Skokie, Illinois). Candidate loxP insertions were identified by co-conversion using a *dpy-10* guide and repair oligo (Arribere *et al*. 2014). All components [pSS4 *dpy-10* guide and Cas9 expression (50 ng/μl), *pbrm-1* guide plasmid (50 ng/μl), loxP repair oligo (30 ng/μl), *dpy-10* repair oligo (30 ng/μl)] were injected into RA650. F1 roller worms were placed three to a plate and allowed to self-fertilize. Once the food was depleted, a portion of the population was washed off the plate and treated with proteinase K to produce a crude DNA prep. These DNA preps were screened using primers in *pbrm-1* and loxP. Individual animals from populations containing a PCR product of the correct size were singled and allowed to give rise to populations with homozygous insertions; the insertion was verified by sequencing; the resulting strain is RA661. Recombination between the loxP sites would be predicted to remove the last six exons of *pbrm-1*, which are shared by *pbrm-1a* and *pbrm-1b,* as well as all GFP coding exons (Fig. 1A).

**Figure 1.**
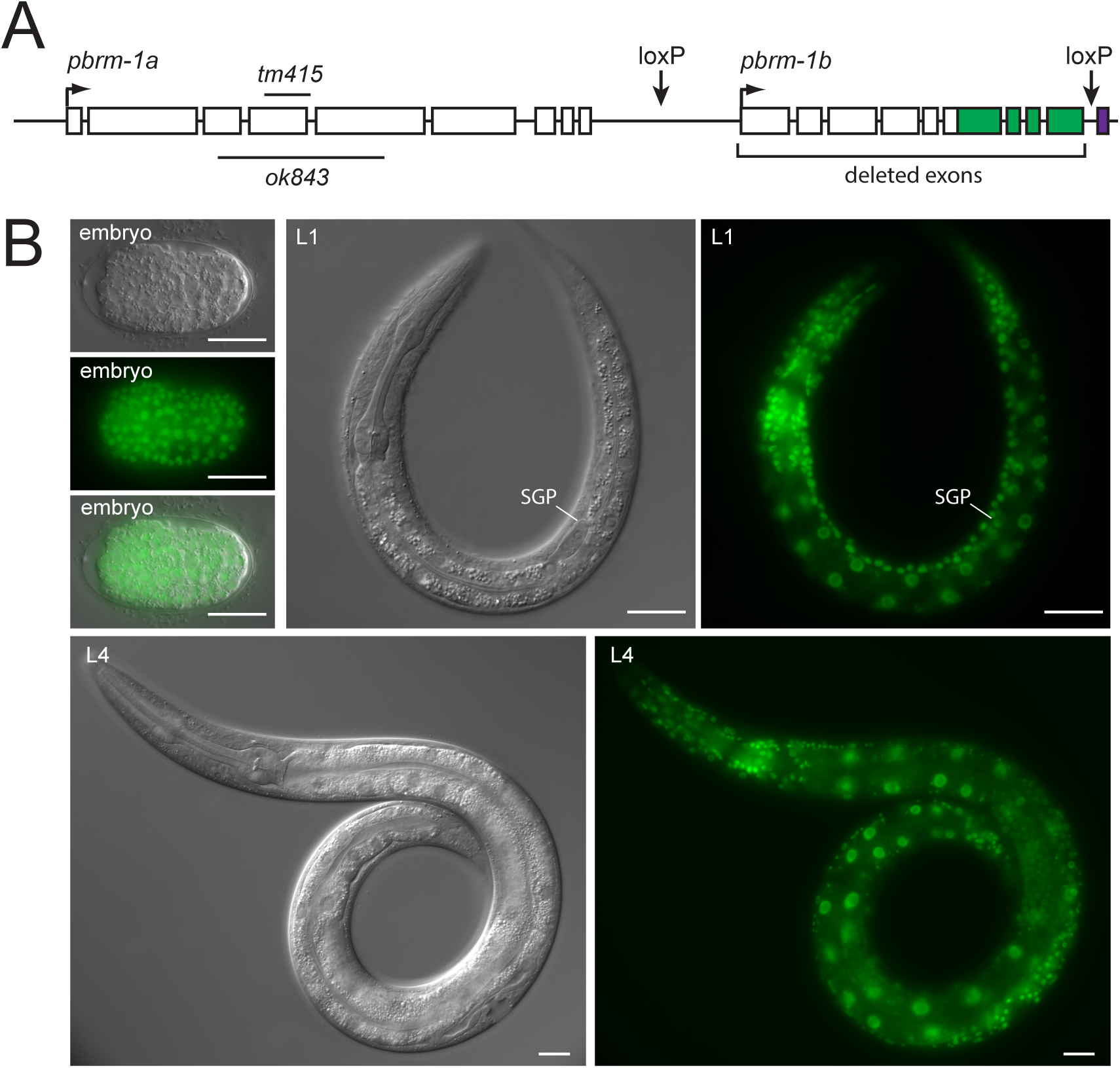
Generation of a GFP-tagged loxP-flanked *pbrm-1* allele. (A) The engineered *pbrm-1* locus contains GFP coding sequences (green exons) downstream of *pbrm-1* coding exons, followed by a 3x FLAG tag (purple). One loxP site remained between GFP and 3x FLAG following removal of the self-excising cassette. An additional loxP site was inserted upstream of the start of *pbrm-1b.* Cre-mediated recombination removes the last six exons of *pbrm-1* and all GFP-encoding exons; the deleted locus is expected to produce a strong loss of function because it removes all common *pbrm-1* exons. (B) PBRM-1::GFP is found in the nucleus of many and perhaps all cells across development, from embryogenesis through adulthood. Images shown are embryos, L1 larvae, and L4 larvae.

### hnd-1 promoter-driven conditional inactivation

*hnd-1p::Cre* (pRA638)*, hnd-1p::GFP-nanobody::ZIF-1::NLS-Cre* (pRA640), and *hnd-1p::ZIF-1* (pRA642) expression constructs were created in pCFJ355 (Addgene #34870), a vector designed for Mos1-mediated single copy transgene insertion on LGX (Frokjaer-Jensen *et al*. 2008; Frokjaer-Jensen *et al*. 2012). The *hnd-1p::Cre* plasmid was generated by digesting pSR47 (Addgene #69258) (Ruijtenberg and van den Heuvel 2015) with PacI and XbaI. The *hnd-1* promoter was amplified by PCR from pJK850 (Mathies *et al*. 2003) using primers RA1396 and RA1397 and inserted in place of the *myo-3* promoter in pSR47. The *hnd-1p::ZIF-1::Cre* and *hnd-1p::GFP-nanobody::ZIF-1::Cre* repair plasmids were generated by digesting pRA638 with PacI and inserting sequences upstream of Cre. ZIF-1 and GFP-nanobody::ZIF-1 sequences were amplified from pOD2046 (Addgene #89367) (Wang *et al*. 2017) using primers RA1506 and RA1507 (GFP-nanobody::ZIF-1) or RA1508 and RA1507 (ZIF-1). The operon linker in pOD2046 was retained to separate the GFP-nanobody::ZIF-1 (or ZIF-1) and Cre coding sequences, creating bicistronic expression constructs. The products were cloned into pRA638 using the NEBuilder HiFi DNA Assembly Master Mix (NEB, Ipswich, MA). All primers are listed in Supplemental Material (File S1, Tables S2).

Single copy insertions were generated using a published protocol (Frokjaer-Jensen *et al*. 2008). Each Cre driver plasmid was injected into *unc-119(ed9); ttTi14024* worms at 50 ng/μl with *eft-3p::transposase* (50 ng/μl) fluorescent co-injection markers [*myo-2p::mCherry* (2.5 ng/μl), *myo-3p::mCherry* (5 ng/μl), and *sur-5p::tdTomato* (5 ng/μl)], and a negative selection marker [*HS::peel-1* (10 ng/μl)]. Resulting non-Unc worms were placed three to a plate and allowed to develop at 25° C until the food was nearly depleted, at which time the worms were moved to 34° C for two hours to eliminate array-bearing animals by *peel-1* negative selection. The plates were screened about one week later for non-Unc animals, which are candidate insertions. Individual animals were singled and their progeny were screened for homozygous insertions. The resulting single copy insertions are *rdIs67 [hnd-1p::Cre]*, *rdIs74 [hnd-1p::GFP-nanobody::ZIF-1::NLS-Cre],* and *rdIs79 [hnd-1p::ZIF-1]*. Each of these transgenes was crossed with RA661 to generate strains RA663 [*pbrm-1(rd31); rdIs67*], RA678 [*pbrm-1(rd31); rdIs74*], and RA685 [*pbrm-1(rd31); rdIs79*]. RA685 was used to control for the effect of the *hnd-1* promoter, *unc-119* rescue, and ZIF-1 protein on gene expression. This allelic combination is referred to as *pbrm-1(control)*.

### Phenotypic analysis

Six first-day adult worms were placed on a plate and allowed to lay eggs for ∼6 hours. Embryos or L1 larvae that remained on the plate after 48 hours were scored as embryonic or L1 lethal, respectively. The L4 staged worms that developed from the embryo collections were examined for gonadogenesis defects using a dissecting microscope. Phenotypes that were classified as gonadogenesis defective were: missing anterior or posterior gonad arm, disorganized gonad with a central patch of gonadal tissue, and gonad absent. Three replicates were performed for each strain and at least 50 worms were scored for each replicate. The percentage of defects across all three replicates is reported in Table 1.

**Table 1.** Phenotypic effects of *pbrm-1* Cre/lox deletion.

**A. Viability**
| Genotype | Lethality (%) |  | n |
| --- | --- | --- | --- |
|  | Embryonic | Larval |  |
| *pbrm-1::GFP(flox)* | 2.6 | 0 | 425 |
| *pbrm-1::GFP(flox); hnd-1p::Cre* | 2.0 | 0 | 200 |
| *pbrm-1::GFP(flox); hnd-1p::GFPnb-ZIF-1::Cre<sup>a</sup>* | 4.7 | 0 | 301 |
| *pbrm-1(rd34)* | 4.4 | 67.7 | 455 |

**B. Gonadogenesis**
| Genotype | Gonad morphology (%) |  |  | n |
| --- | --- | --- | --- | --- |
|  | One arm | Abnormal | Absent |  |
| *pbrm-1::GFP(flox)* | 0.2 | 0 | 0 | 414 |
| *pbrm-1::GFP(flox); hnd-1p::Cre* | 3.1 | 0 | 0 | 196 |
| *pbrm-1::GFP(flox); hnd-1p::GFPnb-ZIF-1::Cre<sup>a</sup>* | 12.5 | 0.3 | 0.3 | 287 |
| *pbrm-1(rd34)* | 20.5 | 4.7 | 3.1 | 127 |

<sup>a</sup> Also called *pbrm-1(TS-KO)*

### RNA sample collection

Five biological replicates were performed on different days and *pbrm-1(control)* and *pbrm-1(TS-KO)* animals were reared and collected in parallel. Populations were grown to adulthood, harvested, and treated with hypochlorite to obtain embryos. Embryos were hatched overnight in sterile M9 medium on a rotating platform to obtain a synchronous population of early L1 larvae. Worms were washed from the plate, purified by sucrose floatation, rinsed once with M9 medium, and stored in Trizol (Ambion, Carlsbad, CA) at -80°C until RNA preparation.

### RNA sequencing and analysis

RNA was isolated using the miRNeasy kit with DNase I digestion performed on the column (Qiagen, Venlo, Netherlands). All samples had RIN numbers of 9.9 or 10 when analyzed using the TapeStation System (Agilent Technologies, Santa Clara, CA). RNA was polyA selected and indexed sequencing libraries were prepared and sequenced by GeneWiz (South Plainfield, NJ). The libraries were sequenced as 150-base, paired-end reads, to an average read depth of 40 million reads per sample using the Illumina HiSeq platform (Illumina, San Diego, CA). The raw RNA sequencing data was examined using FastQC (https://www.bioinformatics.babraham.ac.uk/projects/fastqc/). Sequencing reads were trimmed to remove adapter sequences and low-quality bases. Trimmed reads were mapped to the *C. elegans* genome (Ensembl genome assembly release WBcel325) using the STAR aligner version 2.5.2b (Dobin *et al*. 2013). Unique gene hit counts were calculated using feature Counts from the Subread package v.1.5.2 (Liao *et al*. 2013; Liao *et al*. 2014). Only unique reads that fell within exons were counted. TPM (Transcripts Per Million) values were calculated for each gene. Differential expression was determined using DESeq2 (Love *et al*. 2014). To visualize the variance among replicates and samples, principal component analysis was performed on iDEP (Ge *et al*. 2018) using regularized log transformed data. Volcano plots were generated using R version 4.3.0. Venn diagrams were generated using BioVenn (Hulsen *et al*. 2008).

### Bioinformatics

Overrepresentation of gene ontology (GO) terms for the differentially expressed genes (DEGs) was determined using the statistical overrepresentation test in PANTHER (Thomas *et al*. 2003; Mi *et al*. 2013; Mi *et al*. 2017). Gene lists were compared to all genes with TPM greater than zero in all five replicates of at least one sample type using the GO Biological Process dataset and Fisher’s exact test with false discovery rate (FDR) correction. The DEGs were compared to previously published gene expression datasets from sorted SGPs and hmcs (Mathies *et al*. 2019) and sorted embryonic muscle cells (Fox *et al*. 2007). Gene names were converted to WBGeneIDs using the Gene Name Sanitizer tool on WormBase and lists were compared using the VLOOKUP function in Excel (Microsoft, Redman, WA). The following subsets of *pbrm-1* DEGs were identified: 1-SGP expressed genes (FPKM ≥ 1), 2-Total muscle expressed genes, 3-Muscle enriched genes, 4-SGP-biased genes that had increased expression in *pbrm-1(TS-KO)* (adjusted *p* ≤ 0.05), and 5-hmc-biased genes with reduced expression in *pbrm-1(TS-KO)* (adjusted *p* ≤ 0.05) (File S2).

### Locomotion assays

Ten first day adult worms were placed in copper rings that had been melted into the surface of agar plates. The rings served as corrals to allow for testing of four strains in parallel on one plate. Worms were acclimated to the lack of food for 30 minutes, after which they were moved to test plates. Worm locomotion was recorded for 2 minutes starting at the 10-minute time point. The speed of each worm was calculated using Image Pro Plus software (Media Cybernetics, Inc., Rockville, MD), and an average speed for each group of worms (n = 1) was calculated. Six trials were performed for each genotype; all genotypes were tested simultaneously on the same plates. Two-tailed paired Student’s t tests were used for statistical comparisons of the basal speeds.

#### Reporter validation

Six genes were selected from among the SGP-expressed *pbrm-1(TS-KO)* DEGs for reporter validation; three had higher expression in SGP *vs.* hmc (SGP-biased) and three had higher expression in hmc *vs.* SGP (hmc-biased) in a dataset generated from wild-type SGPs and hmcs (Mathies *et al*. 2019). Genes were prioritized based on highest fold change and lowest adjusted p-values and reporters were made using PCR fusion (Hobert 2002). The pPD95.75 plasmid was modified to use the *tbb-2* 3’ UTR because it promotes high levels of expression (Dour and Nonet 2021) and lacks the background expression reported for the *unc-54* 3’ UTR (Silva-Garcia *et al*. 2019)*. tbb-2* sequence was amplified using primers RA1792 and RA1793 and cloned into pPD95.75 (Addgene plasmid #1494) digested with EcoRI and SpeI using the NEBuilder HiFi DNA Assembly Master Mix (NEB, Ipswich, MA). The resulting plasmid was used as the template for amplification of GFP and 3’ UTR sequences using primers RA1791 and RA1799. Nested forward primers (F1 and F2) and a reverse fusion primer (R) were designed for each gene. Promoter sequences were amplified from genomic DNA using the F1 and R primers and contained up to 5 kb or all sequence to the next upstream gene. The promoter and GFP 3’ UTR PCRs were combined, and a fusion product was amplified using F2 and *tbb-2* 3’ UTR primers. The PCR product was injected into N2 worms at 10-20 ng/μl with 50 ng/μl pRF4 and 50 ng/μl DNA ladder (NEB, Ipswich, MA). The pRF4 plasmid produces a dominant roller phenotype; it was used as a co-injection marker (Mello *et al*. 1991). At least two transmitting lines were isolated for each construct. The line with the highest transmission frequency was crossed into *pbrm-1(ok843)* mutants using *oxTi718* [*eft-3p::tdTomato::H2B*] as a balancer; *oxTi718* is at 2.07 and *pbrm-1* is at 2.10 on LGI. Homozygous *pbrm-1* mutants were identified by the absence tdTomato expression, and their progeny were examined for reporter expression.

Twenty first-day adults were placed on a plate and allowed to lay eggs for two hours. The resulting L1 larvae were examined approximately 16 hours later, using fluorescence and differential interference contrast (DIC) microscopy. Only L1 larvae containing four cells in the gonad primordium were scored because most *pbrm-1* mutants do not develop beyond this stage. GFP fluorescence in SGPs was noted and three levels of expression were recorded – dim, distinct, or bright. At least 30 L1 staged worms were observed. Each SGP was assigned a ranked numerical score for the level of expression (0 = none, 1 = “dim”, 2 = “distinct”, and 3 = “bright”). Statistical comparisons were made using unpaired Student’s t-tests in Prism version 9.5.1 (GraphPad). Reporters were visualized using a Zeiss Axioskop II microscope. All fluorescent images intended for comparison were taken with the same exposure and had identical image adjustments.

## Results

### Conditional inactivation of *pbrm-1* in mesodermal tissues

Strong loss-of-function alleles of *pbrm-1* result in a high degree of embryonic or L1 larval lethality in the progeny of homozygous animals (Large and Mathies 2014). The few escaping animals have incompletely penetrant gonadogenesis defects, indicating that there is a role for *pbrm-1* in the somatic gonad. To facilitate our analysis of this gonadogenesis function, we generated a strain that combines Cre/lox recombination with GFP nanobody-directed protein degradation to eliminate functional PBRM-1 from SGPs, while preserving it in most of the animal. First, we created a *pbrm-1* translational GFP fusion using an established CRISPR/Cas9 genome editing protocol (Dickinson *et al*. 2015). We inserted GFP following the last *pbrm-1* exon to create a C-terminal PBRM-1::GFP fusion protein. Following removal of the selection cassette, this allele retains one loxP site downstream of the GFP coding exons. We inserted a second loxP site upstream of the tenth *pbrm-1* exon, before the start of *pbrm-1b* transcription, creating the loxP-flanked allele (Fig. 1A). This reporter, hereafter called *pbrm-1::GFP(flox),* expresses GFP in most or all cells of the animal, across all life stages (Fig. 1B). The GFP insertion in *pbrm-1* has minimal effects on viability and somatic gonad development (Table 1). When recombined by Cre recombinase, the resulting deletion allele is predicted to lack the last six exons of *pbrm-1*, which are shared by *pbrm-1a* and *pbrm-1b,* as well as the GFP coding sequences, which should cause a loss of function of both *pbrm-1a* and *pbrm-1b*.

To conditionally inactivate *pbrm-1* early in the lives of the SGPs, we used the *hnd-1* promoter to drive a nuclearly-localized Cre recombinase. *hnd-1* is expressed in embryonic mesodermal tissues derived from the MS, C, and D lineages, and it is expressed very early in embryonic SGPs (Mathies *et al*. 2003). We generated single copy insertions of *hnd-1p::Cre* using Mos1-mediated single copy insertion (MosSCI) (Frokjaer-Jensen *et al*. 2008). We found that this driver resulted in Cre-mediated recombination in mesodermal cells, including body wall muscles, the M mesoblast, and SGPs, as assessed using a reporter that switches from *mCherry* to GFP expression upon Cre recombination (Ruijtenberg and van den Heuvel 2015) (Fig. 2A-C). We crossed the Cre driver with *pbrm-1::GFP(flox)* to induce the excision of the *pbrm-1* exons and examined L1 SGPs for expression of PBRM-1::GFP. The *hnd-1p* Cre driver substantially reduced the level of PBRM-1::GFP protein in SGPs, but it did not totally eliminate the protein (Fig 2D-E). The residual GFP fluorescence suggested that PBRM-1 protein was produced from mRNA generated before the excision event and that perdured in SGPs. In order to remove the remaining protein, we employed a GFP nanobody fused to ZIF-1, which targets PBRM-1::GFP for ubiquitin-mediated degradation by the proteosome (DeRenzo *et al*. 2003; Wang *et al*. 2017). We created a bicistronic construct that expresses GFP-nanobody::ZIF-1 and Cre (Fig. 2F). The combination of Cre recombination and ZIF-1-mediated protein degradation eliminated all visible GFP fluorescence from L1 SGPs (Fig. 2F). To control for the effect of ZIF-1 and other genetic elements in the Cre drivers, we generated a construct expressing only ZIF-1, and confirmed that this driver did not affect PBRM-1::GFP protein levels (Fig. 2G).

**Figure 2.**
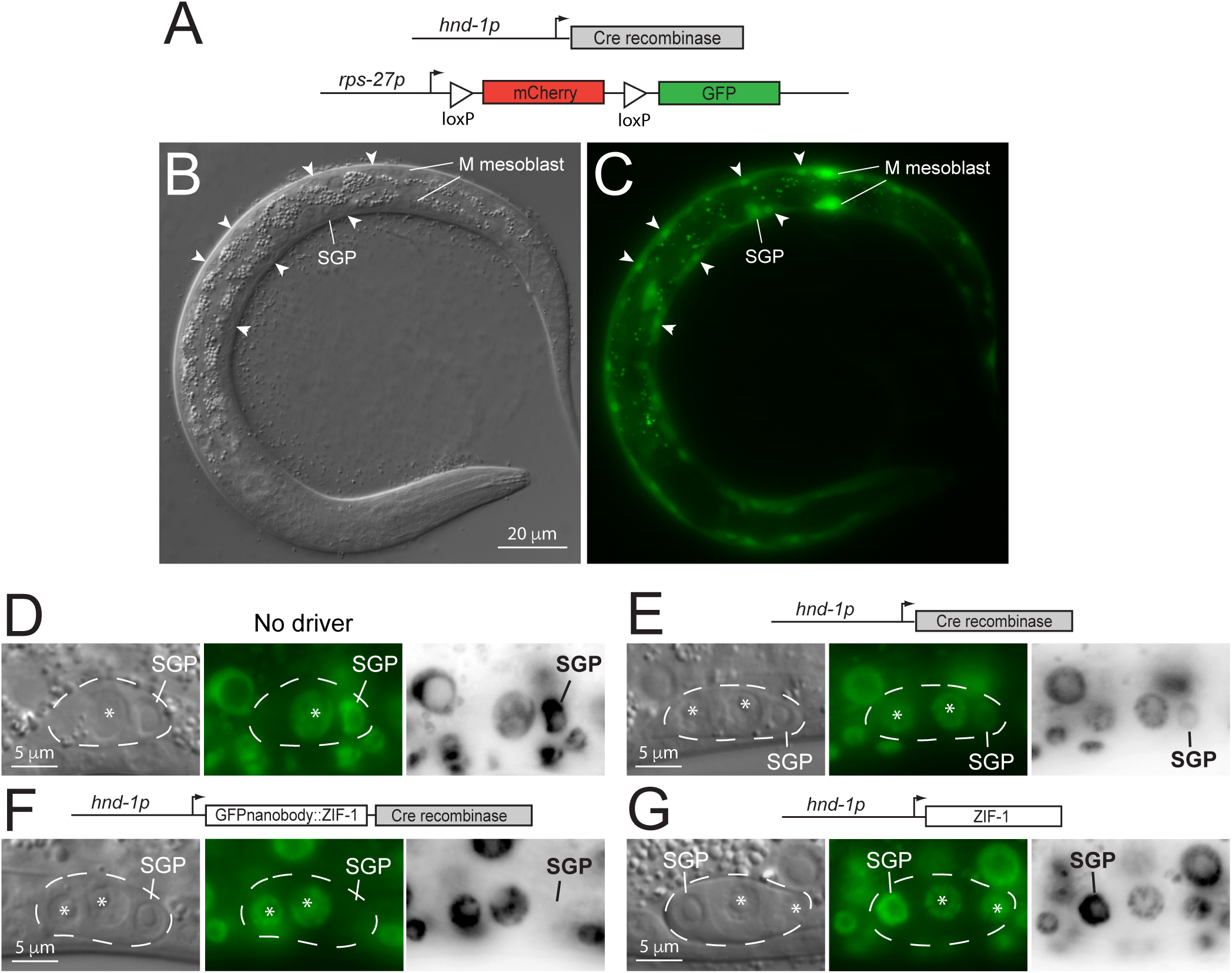
*pbrm-1* conditional knockout in SGPs. (A-C) A Cre readout reporter shows where *hnd-1* promoter driven Cre recombinase is active. The readout reporter is driven by a ubiquitous promoter and switches from mCherry to GFP expression upon recombination between loxP sites. (B-C) *hnd-1p::Cre* promotes recombination in SGPs, body wall muscles (arrow heads), and M mesoblast daughters; DIC image (B) and GFP fluorescence (C). (D-G) A combination of Cre/lox recombination and GFP-nanobody directed protein degradation was used to eliminate PBRM-1::GFP from SGPs. Driver constructs are indicated above images: no driver (D) *hnd-1p::Cre* (E), *hnd-1p::*GFP-nanobody::ZIF-1::Cre (F), *hnd-1p::*ZIF-1 alone (G). DIC image (left), GFP fluorescence (middle), inverted monochrome fluorescent image (right). The gonad primordium (dashed line) and SGPs are indicated, asterisks mark germ cells. All fluorescent images are 1 s exposures with identical adjustments. PBRM-1::GFP fluorescence in SGPs is reduced by *hnd-1p::Cre* recombination (E) and made invisible by the addition of ZIF-1-mediated degradation (F); *hnd-1p::*ZIF-1 does not affect PBRM-1::GFP.

To determine how our tissue-specific inactivation strategies affect *pbrm-1* function, we examined the phenotype of each of the strains (Table 1). *pbrm-1* null or strong loss-of-function alleles result in highly penetrant embryonic or early larval lethality (Large and Mathies 2014). In contrast, all the conditional inactivation strains had minimal lethality (Table 1). Animals lacking *pbrm-1* function maternally and zygotically, and that escape lethality, have incompletely penetrant gonadogenesis defects; typically they are missing one of the two gonad arms (Large and Mathies 2014). We found that *pbrm-1::GFP(flox)* had a low penetrance gonodogenesis defect on its own, *hnd-1p::*Cre-mediated recombination increased this penetrance to 3.1%, and the further addition of GFP nanobody-mediated protein degradation substantially increased the penetrance of the defect to 13.1% (Table 1). Therefore, by employing a combination of Cre/lox recombination and ZIF-1-mediated protein degradation, we have generated a tissue-specific (TS) *pbrm-1* knockout (KO), *pbrm-1(TS-KO),* that produces a strong loss-of-function phenotype in the somatic gonad with minimal lethality and will allow us to perform gene expression studies.

### *pbrm-1-*regulated genes

We used *pbrm-1(TS-KO)* to identify genes that are regulated by *pbrm-1* in mesodermal tissues. The strain containing the ZIF-1 driver (Fig. 2F) served as a control; these worms are *pbrm-1(+)* in all tissues, and they contain the same selectable markers as *pbrm-1(TS-KO);* we refer this strain as *pbrm-1(control).* We previously sorted SGPs for gene expression analysis (Mathies *et al*. 2019). For this study, we chose to isolate mRNA from whole animals because *pbrm-1* loss of function results in SGPs that sometimes fail to express appropriate markers of their fate (Large and Mathies 2014); these SGPs might not be included in the analysis if we sorted based on SGP marker expression. We obtained synchronous populations of early L1 stage worms by bleaching gravid adults and allowing the embryos to hatch in the absence of food. We performed five replicates on different days, and we grew, collected, and processed *pbrm-1(TS-KO)* and *pbrm-1(control)* worms in parallel. RNA sequencing libraries were prepared, sequenced, and mapped to the genome by GeneWiz (Azenta Life Sciences, Plainfield, NJ).

During these experiments, we noticed that some L1 larvae from *pbrm-1(TS-KO)* lacked GFP expression entirely. Because *hnd-1* has a maternal effect (Mathies *et al*. 2003), we thought it was likely that these larvae resulted from maternal recombination of the *pbrm-1* locus. This maternal recombination provided an opportunity to assess the effect of our Cre/lox-mediated deletion on the *pbrm-1* locus. We isolated animals lacking GFP in all tissues, which are homozygous for the *pbrm-1* deletion in the germline, and examined their progeny for lethality and somatic gonad defects. We found that this new deletion, *pbrm-1(rd34),* had phenotypes that were seen in other strong loss-of-function *pbrm-1* alleles (Table 1). When compared to *pbrm-1(ok843)* (Large and Mathies 2014), the *rd34* allele had less lethality and a higher penetrance gonadogenesis defect. Therefore, deletion of the C-terminal *pbrm-1* exons creates a strong loss-of-function allele that is at least as strong as the strongest reported *pbrm-1* allele in the somatic gonad. To determine the prevalence of the maternal *pbrm-1(-)* animals, we examined three different samples and found that 3.1 +/- 1.1 % of the L1 larvae had no GFP fluorescence. Since these L1 larvae are *pbrm-1* mutant in all cells, including SGPs, we reasoned that they should not significantly impact our ability to identify *pbrm-1* regulated genes in SGPs.

We assessed the correlation between biological replicates in our transcriptomic data and found that our sample types were separated by a combination of principal components one and three (Fig. S1). Together, these principal components account for 51% of the variance in the dataset. We examined differential gene expression using DESeq2 (Love *et al*. 2014) and found that 1955 genes were differentially expressed between *pbrm-1(TS-KO)* and *pbrm-1(control)* (FDR ≤ 0.05) (File S2). We did not apply a fold change cutoff to this analysis because we are interested in identifying gene expression changes that occur in two of the 558 cells of the newly hatched L1 larva, and we expect that some of the important changes in SGP expression may not be of large magnitude. Among the differentially expressed genes (DEGs), there were similar numbers of up and down-regulated genes in our dataset (Fig. 3A).

**Figure 3.**
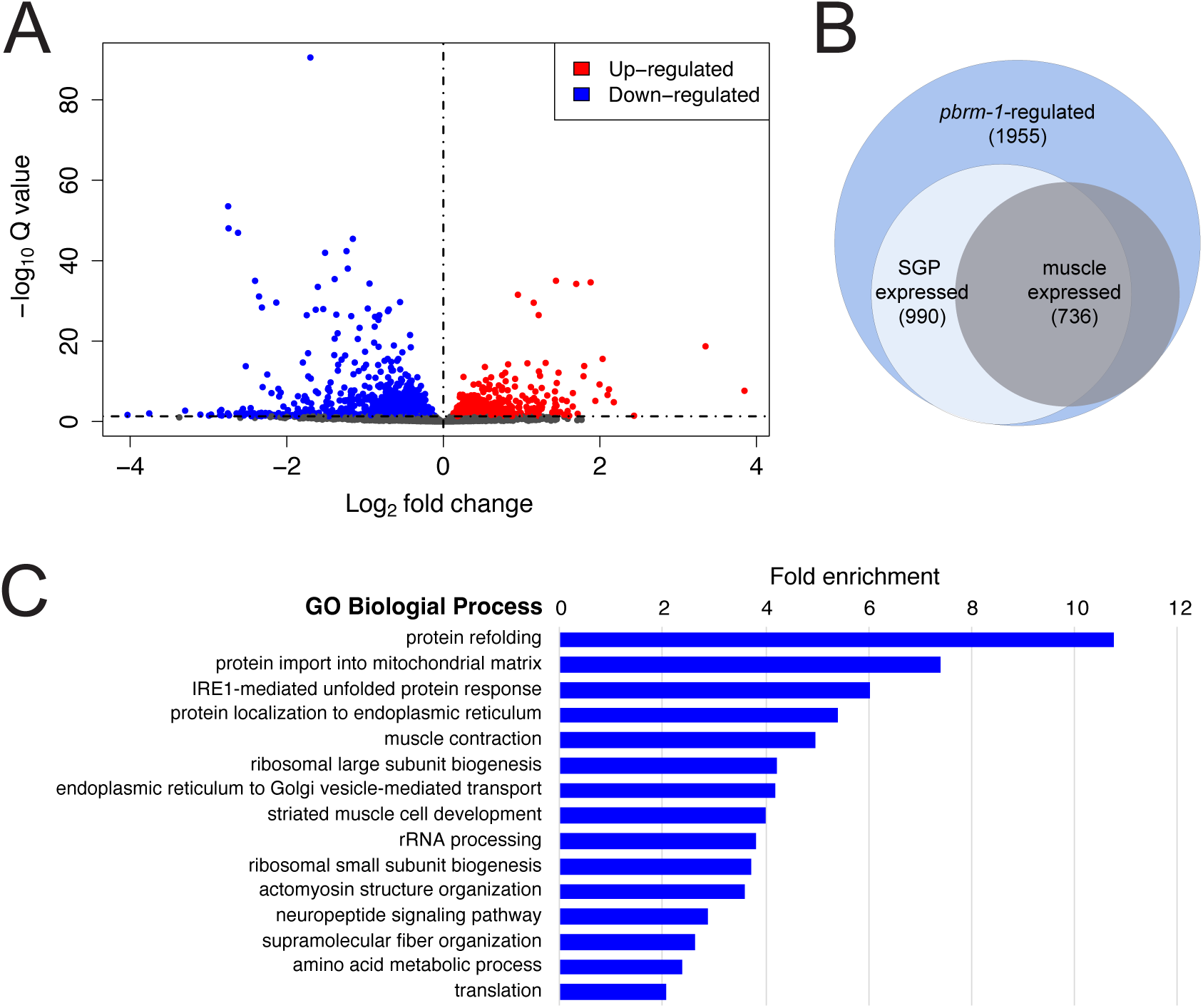
Genes regulated by *pbrm-1* in mesodermal tissues. (A) Volcano plot showing the distribution of differentially expressed genes between *pbrm-1(TS-KO)* and *pbrm-1(control)*. (B) Venn diagram indicating the proportion of *pbrm-1-*regulated genes (dark blue) that are also expressed in SGPs (light blue) or embryonic muscles (gray). The number of genes in each circle is indicated; 558 genes are expressed in both SGPs and muscles (light blue/gray overlap). (C) Gene ontology biological process categories that are statistically over-represented among the SGP-expressed *pbrm-1-*regulated genes. Fold enrichment is plotted.

Our *pbrm-1(TS-KO)* allele removes PBRM-1 from SGPs and other mesodermal cells, primarily body wall muscle (Mathies *et al*. 2003). Therefore, to enrich for genes that function in SGPs, we first filtered our DEGs against all genes that were expressed in SGPs in our transcriptomics analysis of sorted SGPs (Mathies *et al*. 2019). This resulted in 990 candidate *pbrm-1-*regulated genes in SGPs (Fig. 3B, File S2). We examined the genes for Gene Ontology (GO) biological process terms and found two over-represented categories related to muscle function, “muscle contraction” and “striated muscle cell development” (Fig. 3C, File S3). The identification of muscle GO terms suggested that our filter for SGP expressed genes did not eliminate those that also function in muscles. Consistent with this observation, we found that 558 of the 990 *pbrm-1-*regulated genes expressed in SGPs were also expressed in embryonic muscle cells (Fig. 3B; File S2) (Fox *et al*. 2007). Most of the genes in the muscle GO categories were enriched in embryonic muscle cells (Table 2) and nearly all were down-regulated in *pbrm-1(TS-KO),* suggesting that *pbrm-1* promotes muscle differentiation by regulating genes that are essential for muscle function.

**Table 2.** Muscle function genes overrepresented among *pbrm-1* regulated genes.

| **Gene** | **Description** | **Muscle enriched** | ***pbrm-1* expression** |
| --- | --- | --- | --- |
| *dyb-1* | alpha-dystrobrevin | No | Down |
| *lev-11* | tropomyosin | Yes | Down |
| *mel-26* | BTB protein | No | Up |
| *mup-2* | troponin T | Yes | Down |
| *myo-6* | myosin heavy chain | Yes | Down |
| *pat-10* | troponin C | Yes | Down |
| *pat-3* | beta-integrin | Yes | Down |
| *tni-3* | troponin I | Yes | Down |
| *tnt-2* | troponin T | Yes | Down |
| *tnt-4* | troponin T | No | Down |
| *unc-27* | troponin I | Yes | Down |
| *unc-52* | perlecan | Yes | Down |

Since *pbrm-1* regulates the expression of genes required for muscle contraction, we reasoned that we might see defects in locomotion in *pbrm-1(TS-KO)* worms. Locomotion is a neuromuscular process and, mutations affecting muscle function result in uncoordinated movement and reduced locomotion speed (Gieseler *et al*. 2017). Wild-type worms travel at approximately 200 μm/sec in the absence of food in our assays (Table 3). The *pbrm-1* translational GFP insertion did not significantly alter locomotion speed, while both *pbrm-1(TS-KO)* and *pbrm-1(control)* exhibited slower locomotion speeds. Importantly, there was no significant difference in speed between *pbrm-1(TS-KO)* and *pbrm-1(control)*, suggesting that *pbrm-1* is not required for normal locomotion. Both strains are homozygous for a loss-of-function *unc-119* allele, and they carry two wild-type copies of the *C. briggsae unc-119* gene (Frokjaer-Jensen *et al*. 2008). Since *unc-119* mutants have severely reduced locomotion speed (Maduro and Pilgrim 1995), we hypothesize that the selectable *unc-119* marker causes a locomotion defect that might mask any effect of *pbrm-1* on locomotion.

**Table 3.** Locomotion speed of *pbrm-1* strains.

| **Genotype** | **Speed ($\mu\text{m}/\text{sec}$)<sup>a</sup>** | **p-value** |
| --- | --- | --- |
| wild type | 205.9 +/- 3.3 |  |
| *pbrm-1::GFP(flox)* | 199.5 +/- 5.6 | 0.30<sup>b</sup> |
| *pbrm-1(TS-KO)* | 137.8 +/- 9.5 | 0.0023<sup>c</sup>, 0.15<sup>d</sup> |
| *pbrm-1(control)* | 111.8 +/- 7.1 | 0.0004<sup>c</sup> |

<sup>a</sup> Average speed +/- S.E.M.
<sup>b</sup> Compared to wild type
<sup>c</sup> Compared to *pbrm-1::GFP(flox)*
<sup>d</sup> Compared to *pbrm-1(control)*

One other intriguing over-represented GO category is “neuropeptide signaling pathway”. Almost all the *pbrm-1-*regulated genes in this category encode neuropeptide-like proteins or FMRF-like peptides. We previously showed that the sister of the SGPs, the hmc, expresses many secretory proteins, including over 30 FMRF-like peptides (Mathies *et al*. 2019). Because *pbrm-1* is important for the cell fate decision that distinguishes SGPs from their hmc sisters (Large and Mathies 2014), one simple model is that neuropeptide signaling genes are up-regulated in *pbrm-1* mutant SGPs because they are transformed toward the hmc fate. We examined the *pbrm-1-*regulated neuropeptide signaling genes in our SGP/hmc expression dataset and found that six of 22 were normally expressed at higher levels in hmc than in SGPs (hmc-biased), and all of these hmc-biased genes had increased expression in *pbrm-1* mutants (File S3). These genes are excellent candidates for hmc differentiation genes that are up-regulated in SGPs because of the transformation of SGPs to hmcs in *pbrm-1* mutants.

As a first validation step for this dataset, we sought to confirm that *pbrm-1* expression was reduced in *pbrm-1(TS-KO)* when compared with *pbrm-1(control)*. Our differential expression analysis was performed using gene-based counts, and in this analysis, we did not identify *pbrm-1* as a DEG. Cre-mediated recombination of the *pbrm-1* locus only removes exons 10 to 15 (Fig. 1A); therefore, we might not expect to detect differential expression using gene-based counts. We performed differential gene expression analysis at the exon level using DEXSeq (Li *et al*. 2015) and found that five of the last six exons had significantly reduced expression in *pbrm-1(TS-KO)* compared to *pbrm-1(control)* (Fig. S1). We conclude that we can detect *pbrm-1* mRNA expression differences that are restricted to mesodermal tissues in RNA isolated from whole L1 animals.

### *pbrm-1-*regulated genes in SGPs

Our goal with the tissue-specific knockout of *pbrm-1* was to identify genes that are targets of *pbrm-1* in the SGP/hmc fate decision. We are particularly interested in two groups of genes. First are genes that are positively regulated by *pbrm-1* and normally expressed at higher levels in SGPs than in hmc (SGP-biased); we identified 125 genes in this class (File S2); these are candidate positive regulators of the fate and multipotency of SGPs. Second are genes that are negatively regulated by *pbrm-1* and are normally expressed at higher levels in hmc than SGPs (hmc-biased); we identified 53 genes in this class (File S2); these are candidate hmc differentiation genes or negative regulators of multipotency that are repressed in SGPs. We selected three genes in each category for reporter validation: *hsp-4, dpy-18,* and *txdc-1* are normally SGP-biased and are down-regulated in *pbrm-1(TS-KO)*, while *hsp-12.3, acdh-1,* and *nlp-58* are normally hmc-biased and are up-regulated in *pbrm-1(TS-KO)*. We generated transcriptional reporters of each gene and examined them for expression in SGPs or hmc. All the reporters had some expression in L1 larvae. Of the SGP-biased gene reporters, *dpy-18::GFP* and *txdc-1::GFP* had detectable expression in SGPs, while *hsp-4::GFP* did not. Of the hmc-biased genes, only *hsp-12.3::GFP* had detectable expression in hmc.

We chose *hsp-12.3::GFP* and *txdc-12.1::GFP* for further analysis because they had more limited expression in cells other than SGPs and hmc. We crossed each reporter into *pbrm-1(ok843)* mutants and examined their expression patterns. The *hsp-12.3* reporter was expressed in hmc and, only occasionally and very weakly, in SGPs (Fig. 4A-B). In *pbrm-1* mutants, *hsp-12.3::GFP* was expressed more frequently and at higher levels in SGPs (Fig. 4A, D), suggesting that *hsp-12.3* is normally repressed by *pbrm-1* in SGPs. The *txdc-12.1* reporter was expressed in SGPs, and it had similar expression in wild type and *pbrm-1* mutants (Fig. 4B, D), suggesting that either *txdc-12.1* is not regulated by *pbrm-1* in SGPs or our transcriptional reporter does not include all required sequences for proper regulation. Importantly, our reporter analysis identified one target of *pbrm-1* that could play a role in the SGP/hmc cell fate decision, indicating that our data set can be used to identify these genes.

**Figure 4.**
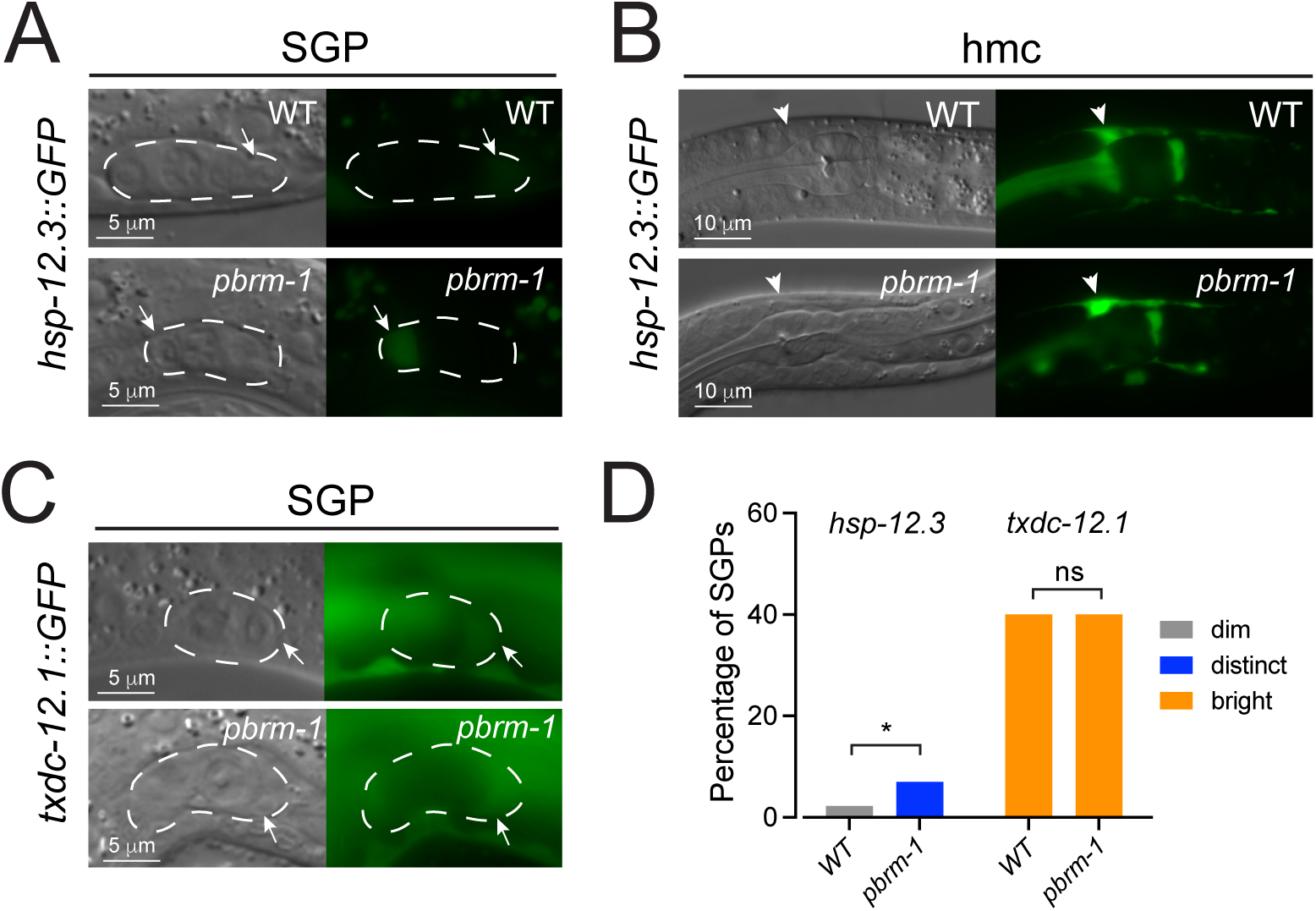
Genes regulated by *pbrm-1* in SGPs. Expression of reporters in wild type (WT) and *pbrm-1(ok843)* mutants. Paired images are DIC (left) and GFP fluorescence (right). Arrows point to SGPs; arrowheads point to hmc; the gonad primordium is outlined (dashed line). Scale bars are indicated. (A-B) *hsp-12.3::GFP* expression in SGP (A) and hmc (B); fluorescent exposures are 500 ms. (A) *hsp-12.3::GFP* is expressed very faintly in wild type SGPs (top) and this expression increases in *pbrm-1(ok843)* mutants (bottom). (B) *hsp-12.3::GFP* is expressed in hmc (top); this expression is unchanged in *pbrm-1(ok843)* mutants (bottom). (C) *txdc-12.1::GFP* is expressed in wild type SGPs (top) and this expression is unchanged in *pbrm-1(ok843)* mutants (bottom); fluorescent exposures are 50 ms. Paired wild type and *pbrm-1* mutant images were taken with identical exposures and adjustments for comparison. (D) Percentage of SGPs with dim (gray), distinct (blue), or bright (orange) GFP in SGPs. Unpaired Student’s t-tests were used to compare expression in wild type and *pbrm-1* mutants; * p ≤ 0.05.

## Discussion

*pbrm-1* encodes the signature subunit of the PBAF complex, which is important for distinguishing multipotent SGPs from their differentiated sister cell, the hmc (Large and Mathies 2014). We used a tissue-specific gene inactivation strategy to identify genes that are regulated by *pbrm-1* in SGPs. Our approach used Cre/lox recombination to remove *pbrm-1* exons that are common to all PBRM-1 isoforms and GFP nanobody-targeted protein degradation to remove residual protein. This combination produced a strong loss-of-function phenotype in the somatic gonad and the absence of any visible PBRM-1::GFP in SGPs, strongly suggesting that PBRM-1 targets in SGPs are among the differentially expressed genes in *pbrm-1(TS-KO)*.

Cre/lox recombination of our floxed *pbrm-1* allele results in the deletion of PBRM-1B and truncation of PBRM-1A after amino acid 1346, removing the DNA binding HMG domain. The strongest existing *pbrm-1* allele by phenotypic and molecular criteria is *ok843,* which truncates PBRM-1A after amino acid 390 and leaves PBRM-1B unaffected (Fig. 1A) (Large and Mathies 2014). Animals carrying the Cre/lox deletion, *rd34*, in their germline had phenotypes similar to *pbrm-1(ok843)*, including embryonic and larval lethality and missing gonadal arms, indicating that this deletion causes a strong loss of gene function. The *rd34* allele resulted in less lethality and a higher penetrance gonadogenesis defect than *ok843*, suggesting that different PBRM-1 domains may be important for these two developmental functions of the gene. The stronger effect of *pbrm-1(rd34)* on gonadogenesis could point to important functions for PBRM-1B in the somatic gonad. Conversely, the stronger effect of *pbrm-1(ok843)* on viability suggests that protein domains removed by this deletion may be important for embryogenesis or early larval development.

Cre/lox deletion of the last six exons of *pbrm-1* produced a phenotype that resembled zygotic loss of function for strong *pbrm-1* deletion alleles in the somatic gonad. *pbrm-1* alleles exhibit maternal effects on somatic gonad development (Large and Mathies 2014), which can be explained by inheritance of maternal PBRM-1 protein by SGPs. Consistent with this, the addition of PBRM-1 degradation in the mesoderm produced a phenotype that resembled maternal and zygotic loss of *pbrm-1* function in the somatic gonad. Our observations are consistent with a previous study that showed dose-dependent functions of the SWI/SNF complex in the regulation of cell division in the M mesoblast: incomplete loss of *swsn-1* via Cre/lox recombination or GFP-nanobody-directed protein degradation produced an over-proliferation phenotype, while the combination of both produced an under-proliferation phenotype (van der Vaart *et al*. 2020). Together, these two studies argue strongly for the use of both genetic deletion and protein degradation to produce a strong loss of gene function in specific tissues.

### *pbrm-1* regulated genes in the SGP/hmc fate decision

To understand how *pbrm-1* influences the SGP/hmc fate decision, we sought to identify genes whose regulation and expression suggested a role in this cell fate decision. We identified genes in two categories: 1-those that were positively regulated by *pbrm-1* and normally SGP-biased (125 genes) and 2-those that were negatively regulated by *pbrm-1* and normally hmc-biased (53 genes). We used fluorescent reporters to examine the expression and regulation of three genes in each category. Out of the six reporters, three had the expected expression in SGPs or hmc. This is consistent with our previous work, in which we found that two of five genes with expression in sorted SGPs were validated by reporters (Mathies *et al*. 2019). There are many reasons that traditional transgenic reporters may not accurately reflect the expression of the gene, the most significant of which is that they do not contain all relevant regulatory sequences. Indeed, in our previous work we showed that, when we used CRISPR/Cas9 genome editing to make an endogenous reporter for a gene that did not mirror our RNA sequencing results, we observed the expected expression pattern, strongly supporting the notion that transcriptional reporters may not always accurately recapitulate the endogenous expression pattern.

We further examined the regulation of two reporters, *hsp-12.3* and *txdc-12.1. hsp-12.3* was up-regulated in *pbrm-1(TS-KO)* and, consistent with this, we found that the reporter was more highly expressed in SGPs in *pbrm-1* mutants, indicating that *hsp-12.3* is normally repressed by *pbrm-1* in SGPs. *txdc-12.1* was down-regulated in *pbrm-1(TS-KO)*, but the reporter did not show any change in expression in *pbrm-1* mutant SGPs. One explanation for this result is that *txdc-12.1* is not regulated by *pbrm-1* in SGPs; instead, it may be regulated by *pbrm-1* in other mesodermal tissues. Alternatively, our reporter might be missing the regulatory sequences that mediate regulation by *pbrm-1* in SGPs. Ultimately, it will be necessary to examine reporters in the native genomic context to fully characterize the regulation of these genes by *pbrm-1*.

Our tissue-specific inactivation strategy provides a significant improvement over gene expression studies using conventional germline mutations, which affect all cells in the animal and that result in a high degree of lethality. However, our approach does have some limitations for identifying *pbrm-1-*regulated genes in SGPs. First, to inactivate *pbrm-1* early in the SGP lineage, we also had to inactivate it in other mesodermal tissues. The *hnd-1* promoter is, to our knowledge, the earliest acting promoter in SGPs; it is expressed shortly after the SGPs are born and while they are migrating to join the PGCs and form the gonad primordium (Mathies *et al*. 2003). The *hnd-1* promoter also drives expression earlier in the MS, C, and D lineages, which produce a total of 83 cells including 70 body muscles, two SGPs, the hmc, and ten other mesodermally-derived cells (Sulston *et al*. 1983). We therefore expect our dataset to include *pbrm-1-*regulated genes in each of these cell types. Since there are many more muscle cells than SGPs, we anticipated that we would identify *pbrm-1-*regulated genes in muscles and, indeed, GO terms related to muscle function were an over-represented category in our DEGs. Second, our *hnd-1* driven Cre recombinase causes a low level of maternal recombination resulting in animals that are mutant for *pbrm-1* in all 558 cells of the L1 larva, including SGPs. Both limitations will reduce our sensitivity to detect *pbrm-1-*regulated genes in SGPs. Nonetheless, we identified 178 candidate genes and confirmed one gene, *hsp-12.3,* that is regulated by *pbrm-1* in SGPs.

### Muscle differentiation genes are positively regulated by *pbrm-1*

*hnd-*1-driven gene inactivation eliminates PBRM-1 protein from most of the body wall muscles. We found that genes with GO terms related to muscle function were over-represented among the *pbrm-1-*regulated genes in our dataset. The *C. elegans* body wall musculature differentiates in embryogenesis just prior to the two-fold stage (Hresko *et al*. 1994). Our gene expression analysis was performed on L1 stage larvae, in which muscles are fully differentiated. We found 12 DEGs related to muscle contraction or striated muscle development, and all but one of these genes had reduced expression in *pbrm-1(TS-KO),* indicating that *pbrm-1* normally promotes the expression of these genes.

We predicted that the loss of muscle gene expression might affect worm locomotion. However, we did not observe any difference in locomotion speed between *pbrm-1(-)* and *pbrm-1(+)* worms, suggesting that *pbrm-1* is not required in muscles for normal locomotion. One possibility is that the gene expression changes were not significant enough to impact muscle structure and therefore affect locomotion. Most of the muscle genes had modestly reduced expression, ranging from 70-90% of the levels in *pbrm-1(+)* (File S3). This analysis was complicated by the fact that both strains used for our RNA-Seq analysis had reduced locomotion speed compared to wild type. These strains carry single copy insertions that were generated using the MosSCI technique, which uses rescue of an *unc-119* mutant phenotype as a selectable marker (Frokjaer-Jensen *et al*. 2008). *unc-119* mutants have significantly reduced locomotion speed (Maduro and Pilgrim 1995). Therefore, it is likely that the reduced locomotion speed in these strains is due to incomplete rescue of the *unc-119* phenotype and, it is possible that the locomotion defect may be masking any subtle effect of mesodermal *pbrm-1* inactivation on locomotion.

Mammalian SWI/SNF complexes promote MyoD-dependent muscle differentiation, and about a third of MyoD-induced genes require SWI/SNF function (de la Serna *et al*. 2001; de la Serna *et al*. 2005). These studies specifically implicated the core ATPase subunits of SWI/SNF, brahma (Brm) and brahma-related gene-1 (Brg1). *C. elegans* has a single gene, *swsn-4,* encoding the SWI/SNF ATPase subunit (Sawa *et al*. 2000). Based on phenotypic analyses, it is predicted that SWSN-4 is incorporated into both BAF and PBAF complexes (Shibata *et al*. 2012; Large and Mathies 2014). Our finding that PBRM-1 regulates the expression of muscle differentiation genes in *C. elegans* raises the possibility that the PBAF complex promotes muscle differentiation across phyla. It further suggests that our dataset may provide insight into how PBAF regulates the differentiation of muscles.

## Data Availability

Strains are available upon request. The RNA sequencing dataset generated during this study is available in the NCBI SRA repository, accession number PRJNA1027254.

## Acknowledgements

Strains were provided by the *Caenorhabditis* Genetics Center.

---

**Figure S1.**
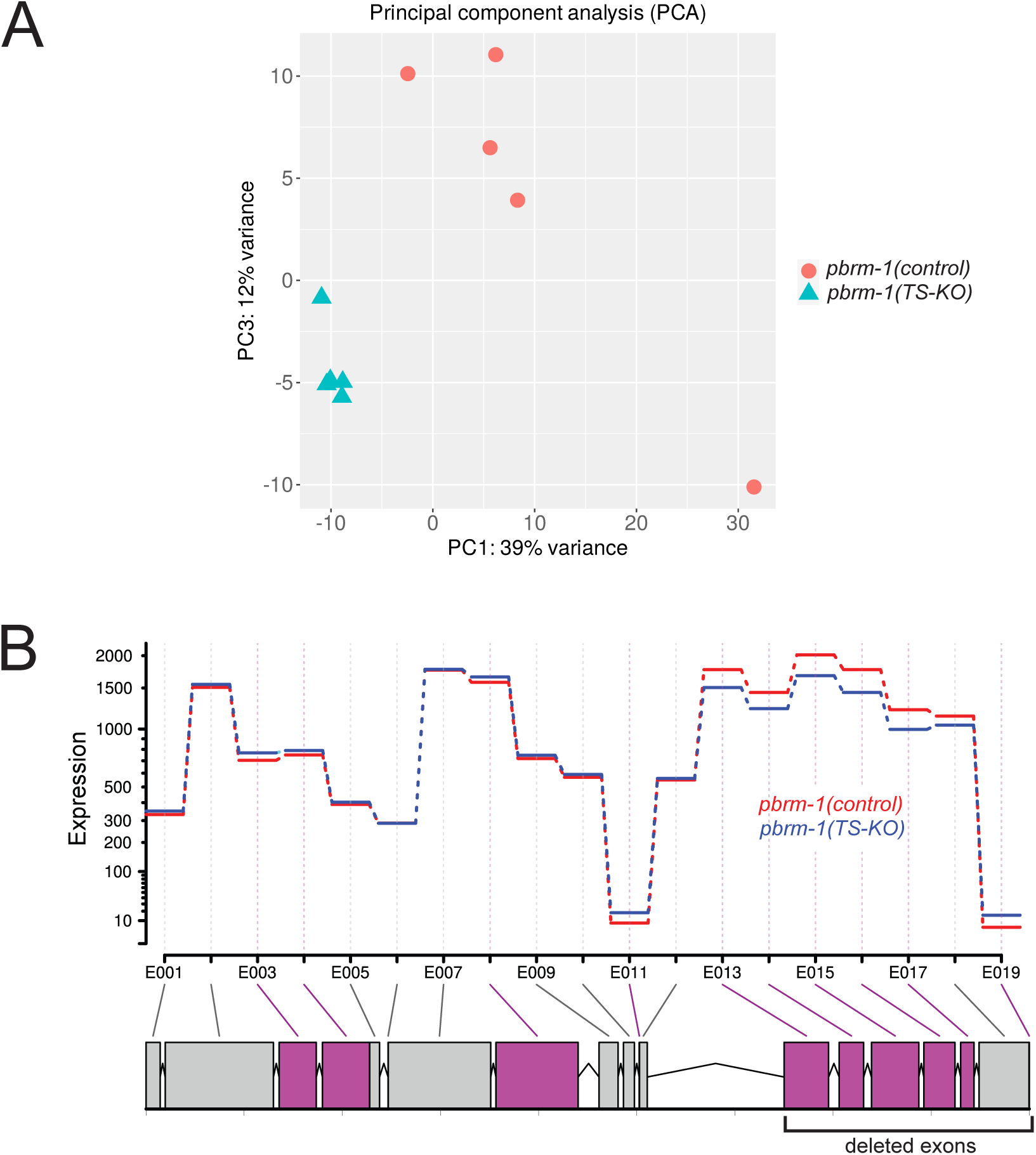
RNA sequencing analysis. (A) Principal component analysis. Gene expression profiles plotted against principal components one and three (PC1 and PC3). *pbrm-1(TS-KO)* and *pbrm-1(control)* replicates group separately. (B) Exon-level differential gene expression analysis for *pbrm-1.* Expression level in *pbrm-1(TS-KO)* (blue) or *pbrm-1(control)* (red) is plotted for each exon. Significant differences are indicated by pink exons on the gene diagram. Exons deleted by Cre/lox recombination in the *pbrm-1(TS-KO)* allele are indicated. E006, E011, and E019 do not correspond to known exons and are only 2 or 3 bps in length.

## References

Arribere JA, Bell RT, Fu BX, Artiles KL, Hartman PS, Fire AZ. 2014. Efficient marker-free recovery of custom genetic modifications with CRISPR/Cas9 in *Caenorhabditis elegans*. Genetics 198: 837–846. 10.1534/genetics.114.169730

Brenner S. 1974. The genetics of *Caenorhabditis elegans*. Genetics 77: 71–94.

Clapier CR, Cairns BR. 2009. The biology of chromatin remodeling complexes. Annu Rev Biochem 78: 273–304. 10.1146/annurev.biochem.77.062706.153223

de la Serna IL, Carlson KA, Imbalzano AN. 2001. Mammalian SWI/SNF complexes promote MyoD-mediated muscle differentiation. Nat Genet 27: 187–190. 10.1038/84826

de la Serna IL, Ohkawa Y, Berkes CA, Bergstrom DA, Dacwag CS, Tapscott SJ, Imbalzano AN. 2005. MyoD targets chromatin remodeling complexes to the myogenin locus prior to forming a stable DNA-bound complex. Mol Cell Biol 25: 3997–4009. 10.1128/MCB.25.10.3997-4009.2005

DeRenzo C, Reese KJ, Seydoux G. 2003. Exclusion of germ plasm proteins from somatic lineages by cullin-dependent degradation. Nature 424: 685–689. 10.1038/nature01887

Dickinson DJ, Pani AM, Heppert JK, Higgins CD, Goldstein B. 2015. Streamlined Genome Engineering with a Self-Excising Drug Selection Cassette. Genetics 200: 1035–1049. 10.1534/genetics.115.178335

Dobin A, Davis CA, Schlesinger F, Drenkow J, Zaleski C, Jha S, Batut P, Chaisson M, Gingeras TR. 2013. STAR: ultrafast universal RNA-seq aligner. Bioinformatics 29: 15–21. 10.1093/bioinformatics/bts635

Dour S, Nonet M. 2021. Optimizing expression of a single copy transgene in *C. elegans*. MicroPubl Biol 2021. 10.17912/micropub.biology.000394

Fox RM, Watson JD, Von Stetina SE, McDermott J, Brodigan TM, Fukushige T, Krause M, Miller DM, 3rd. 2007. The embryonic muscle transcriptome of *Caenorhabditis elegans*. Genome biology 8: R188.

Friedland AE, Tzur YB, Esvelt KM, Colaiacovo MP, Church GM, Calarco JA. 2013. Heritable genome editing in C. elegans via a CRISPR-Cas9 system. Nat Methods 10: 741–743. 10.1038/nmeth.2532

Frokjaer-Jensen C, Davis MW, Ailion M, Jorgensen EM. 2012. Improved Mos1-mediated transgenesis in C. elegans. Nat Methods 9: 117–118. 10.1038/nmeth.1865

Frokjaer-Jensen C, Davis MW, Hopkins CE, Newman BJ, Thummel JM, Olesen SP, Grunnet M, Jorgensen EM. 2008. Single-copy insertion of transgenes in *Caenorhabditis elegans*. Nat Genet 40: 1375–1383.

Ge SX, Son EW, Yao R. 2018. iDEP: an integrated web application for differential expression and pathway analysis of RNA-Seq data. BMC bioinformatics 19: 534.

Gieseler K, Qadota H, Benian GM. 2017. Development, structure, and maintenance of *C. elegans* body wall muscle. in WormBook, pp. 1–59.

Ho L, Ronan JL, Wu J, Staahl BT, Chen L, Kuo A, Lessard J, Nesvizhskii AI, Ranish J, Crabtree GR. 2009. An embryonic stem cell chromatin remodeling complex, esBAF, is essential for embryonic stem cell self-renewal and pluripotency. Proc Natl Acad Sci U S A 106: 5181–5186. 10.1073/pnas.0812889106

Hobert O. 2002. PCR fusion-based approach to create reporter gene constructs for expression analysis in transgenic C. elegans. BioTechniques 32: 728–730.

Hresko MC, Williams BD, Waterston RH. 1994. Assembly of body wall muscle and muscle cell attachment structures in Caenorhabditis elegans. J Cell Biol 124: 491–506. 10.1083/jcb.124.4.491

Hulsen T, de Vlieg J, Alkema W. 2008. BioVenn - a web application for the comparison and visualization of biological lists using area-proportional Venn diagrams. BMC Genomics 9: 488. 10.1186/1471-2164-9-488

Kimble J, Hirsh D. 1979. The postembryonic cell lineages of the hermaphrodite and male gonads in *Caenorhabditis elegans*. Dev Biol 70: 396–417.

Large EE, Mathies LD. 2014. *Caenorhabditis elegans* SWI/SNF subunits control sequential developmental stages in the somatic gonad. G3 (Bethesda) 4: 471–483. 10.1534/g3.113.009852

Lessard J, Wu JI, Ranish JA, Wan M, Winslow MM, Staahl BT, Wu H, Aebersold R, Graef IA, Crabtree GR. 2007. An essential switch in subunit composition of a chromatin remodeling complex during neural development. Neuron 55: 201–215. 10.1016/j.neuron.2007.06.019

Lessard JA, Crabtree GR. 2010. Chromatin regulatory mechanisms in pluripotency. Annu Rev Cell Dev Biol 26: 503–532. 10.1146/annurev-cellbio-051809-102012

Li Y, Rao X, Mattox WW, Amos CI, Liu B. 2015. RNA-Seq Analysis of Differential Splice Junction Usage and Intron Retentions by DEXSeq. PLoS One 10: e0136653. 10.1371/journal.pone.0136653

Liao Y, Smyth GK, Shi W. 2013. The Subread aligner: fast, accurate and scalable read mapping by seed-and-vote. Nucleic Acids Res 41: e108. 10.1093/nar/gkt214

Liao Y, Smyth GK, Shi W. 2014. featureCounts: an efficient general purpose program for assigning sequence reads to genomic features. Bioinformatics 30: 923–930. 10.1093/bioinformatics/btt656

Love MI, Huber W, Anders S. 2014. Moderated estimation of fold change and dispersion for RNA-seq data with DESeq2. Genome Biol 15: 550. 10.1186/s13059-014-0550-8

Maduro M, Pilgrim D. 1995. Identification and cloning of *unc-119*, a gene expressed in the *Caenorhabditis elegans* nervous system. Genetics 141: 977–988.

Mathies LD, Henderson ST, Kimble J. 2003. The *C. elegans* Hand gene controls embryogenesis and early gonadogenesis. Development 130: 2881–2892.

Mathies LD, Ray S, Lopez-Alvillar K, Arbeitman MN, Davies AG, Bettinger JC. 2019. mRNA profiling reveals significant transcriptional differences between a multipotent progenitor and its differentiated sister. BMC Genomics 20: 427. 10.1186/s12864-019-5821-z

Mello CC, Kramer JM, Stinchcomb D, Ambros V. 1991. Efficient gene transfer in *C. elegans*: extrachromosomal maintenance and integration of transforming sequences. EMBO J 10: 3959–3970.

Mi H, Huang X, Muruganujan A, Tang H, Mills C, Kang D, Thomas PD. 2017. PANTHER version 11: expanded annotation data from Gene Ontology and Reactome pathways, and data analysis tool enhancements. Nucleic Acids Res 45: D183–D189. 10.1093/nar/gkw1138

Mi H, Muruganujan A, Casagrande JT, Thomas PD. 2013. Large-scale gene function analysis with the PANTHER classification system. Nat Protoc 8: 1551–1566. 10.1038/nprot.2013.092

Ruijtenberg S, van den Heuvel S. 2015. G1/S Inhibitors and the SWI/SNF Complex Control Cell-Cycle Exit during Muscle Differentiation. Cell 162: 300–313. 10.1016/j.cell.2015.06.013

Sawa H, Kouike H, Okano H. 2000. Components of the SWI/SNF complex are required for asymmetric cell division in *C. elegans*. Mol Cell 6: 617–624.

Shibata Y, Uchida M, Takeshita H, Nishiwaki K, Sawa H. 2012. Multiple functions of PBRM-1/Polybromo- and LET-526/Osa-containing chromatin remodeling complexes in *C. elegans* development. Developmental biology 361: 349–357. 10.1016/j.ydbio.2011.10.035

Silva-Garcia CG, Lanjuin A, Heintz C, Dutta S, Clark NM, Mair WB. 2019. Single-Copy Knock-In Loci for Defined Gene Expression in *Caenorhabditis elegans*. G3 (Bethesda) 9: 2195–2198. 10.1534/g3.119.400314

Singhal N, Graumann J, Wu G, Arauzo-Bravo MJ, Han DW, Greber B, Gentile L, Mann M, Scholer HR. 2010. Chromatin-Remodeling Components of the BAF Complex Facilitate Reprogramming. Cell 141: 943–955. 10.1016/j.cell.2010.04.037

Sulston JE, Schierenberg E, White JG, Thomson JN. 1983. The embryonic cell lineage of the nematode *Caenorhabditis elegans*. Dev Biol 100: 64–119.

Thomas PD, Kejariwal A, Campbell MJ, Mi H, Diemer K, Guo N, Ladunga I, Ulitsky-Lazareva B, Muruganujan A, Rabkin S et al. 2003. PANTHER: a browsable database of gene products organized by biological function, using curated protein family and subfamily classification. Nucleic Acids Res 31: 334–341.

van der Vaart A, Godfrey M, Portegijs V, van den Heuvel S. 2020. Dose-dependent functions of SWI/SNF BAF in permitting and inhibiting cell proliferation in vivo. Sci Adv 6: eaay3823. 10.1126/sciadv.aay3823

Wang S, Tang NH, Lara-Gonzalez P, Zhao Z, Cheerambathur DK, Prevo B, Chisholm AD, Desai A, Oegema K. 2017. A toolkit for GFP-mediated tissue-specific protein degradation in C. elegans. Development 144: 2694–2701. 10.1242/dev.150094

Wood WB. 1988. Introduction to *C. elegans* Biology. in The Nematode Caenorhabditis elegans (ed. WB Wood), pp. 1–16. Cold Spring Harbor Laboratory Press, Cold Spring Harbor, NY.

Wu JI, Lessard J, Crabtree GR. 2009. Understanding the words of chromatin regulation. Cell 136: 200–206. 10.1016/j.cell.2009.01.009

